# Proteomic survey of the DNA damage response in *Caulobacter crescentus*

**DOI:** 10.1101/2023.03.24.534141

**Authors:** Tommy F. Tashjian, Rilee D. Zeinert, Stephen J. Eyles, Peter Chien

## Abstract

The bacterial DNA damage response is a critical, coordinated response to endogenous and exogenous sources of DNA damage. Response dynamics are dependent on coordinated synthesis and loss of relevant proteins. While much is known about its global transcriptional control, changes in protein abundance that occur upon DNA damage are less well characterized at the system level. Here, we perform a proteome-wide survey of the DNA damage response in *Caulobacter crescentus*. We find that while most protein abundance changes upon DNA damage are readily explained by changes in transcription, there are exceptions. The survey also allowed us to identify the novel DNA damage response factor, YaaA, which has been overlooked by previously published, transcription-focused studies.

A similar survey in a Δ*lon* strain was performed to explore *lon’s* role in DNA damage survival. The Δ*lon* strain had a smaller dynamic range of protein abundance changes in general upon DNA damage compared to the wild type strain. This system-wide change to the dynamics of the response may explain this strain’s sensitivity to DNA damage. Our proteome survey of the DNA damage response provides additional insight into the complex regulation of stress response and nominates a novel response factor that was overlooked in prior studies.

**IMPORTANCE:** The DNA damage response helps bacteria to react to and potentially survive DNA damage. The mutagenesis induced during this stress response contributes to the development of antibiotic resistance. Understanding how bacteria coordinate their response to DNA damage could help us to combat this growing threat to human health. While the transcriptional regulation of the bacterial DNA damage response has been characterized, this study is the first to our knowledge to assess the proteomic response to DNA damage in *Caulobacter*.

## INTRODUCTION

The bacterial DNA damage response is a major driver of antibiotic resistance development (1, 2) and the horizontal gene transfer that spreads resistance and pathogenicity from cell to cell (3–6). The canonical DNA damage response was first characterized in *Escherichia coli*. During DNA damage ssDNA accumulates in the cell and is bound by the cell’s main recombinase RecA to form a nucleoprotein filament that induces the autocleavage of the global transcriptional repressor LexA (7, 8). Cleavage of LexA allows transcription of more than 40 genes related to DNA damage tolerance and repair, including *recA* and *lexA* themselves (9, 10). The energy-dependent protease ClpXP then degrades the DNA binding N-terminal domain of LexA to prevent residual repression (11).

As DNA damage is repaired, accumulation of uncleaved LexA represses transcription of DNA damage response genes. Shutdown of the response is also aided by proteolysis of certain response factors (e.g. UmuD’ (12) and UvrA (13) by ClpXP and UmuD (14) and SulA (15) by Lon). In fact, both ClpXP (11) and Lon (16, 17) contribute to the survival of DNA damage in *E. coli*.

The LexA/RecA-based transcriptional control of the response is conserved in *Caulobacter crescentus*, as are many of the genes in the *E. coli* LexA regulon (18). A second, LexA-independent DNA damage response pathway that is dependent on global transcriptional regulator DriD has also been identified and characterized in *Caulobacter* (19, 20). While the energy-dependent protease Lon is conserved in *Caulobacter* and contributes to survival of DNA damage (21), Lon’s confirmed DNA damage-related substrates (UmuD (14) and SulA (15)) are not. Lon has been shown to contribute to DNA damage survival in *Caulobacter* but its role in response to DNA damage remains unclear.

Transcriptome measurements upon treatment with various DNA damaging agents and a bioinformatic search for LexA binding sites have nominated a set of genes that are upregulated upon damage in *Caulobacter* (18, 19, 22). While transcription and proteolysis both play roles in DNA damage response induction and shut-off, relatively little is known about how well transcriptional changes correspond to protein abundance changes at the proteome level.

Here, we present a proteomic survey to investigate the dynamics of protein abundance during the *Caulobacter* DNA damage response. First, we measured protein abundance changes upon DNA damage treatment. Next, we measured changes following translation shutoff to characterize potential proteolysis substrates. We found that while most changes in protein abundance upon DNA damage are explained by changes in transcription, there are exceptions to this rule. By combining our proteomics and previously measured changes in transcript levels upon mitomycin C (MMC) treatment (19), we were able to identify the novel DNA damage response factor YaaA, which has previously been overlooked in surveys that considered transcriptional changes alone. Our validation experiments showed this factor to be important during DNA damage, demonstrating the utility of our approach.

In the second stage of this study, we attempted to identify novel Lon substrates to explain the importance of this protease to DNA damage survival. While we were unable to identify any novel Lon substrates that sufficiently explain this phenotype, we did characterize system-wide changes to the DNA damage response in a Δ*lon* strain. The magnitude of protein abundance changes upon DNA damage are dampened in the Δ*lon* strain compared to those in the wild type strain. This effect can be attributed in part to the elevated basal levels of DNA damage proteins in the untreated Δ*lon* strain (23). This suggests Lon may be of wider importance to the regulation of the DNA damage response than simply degrading of one or a few key substrates.

This study provides new insights into Lon’s broader importance to protein dynamics in *Caulobacter’s* DNA damage response and identifies a novel factor that is important for survival to DNA damage.

## RESULTS

### Proteomics yields novel insights into the DNA damage response

To characterize protein abundance changes during the DNA damage response, we first used the sentinel protein RecA to determine the timescale of an acute MMC treatment in *Caulobacter* (Figure 1A and S1). Cells were treated with MMC for 1 hour and RecA levels were monitored through recovery (Figure 1A and S1). RecA levels peaked 3 hours after the addition of MMC (Figure 1A and S1), which is similar to what was previously observed during chronic MMC treatment (24).

**Figure 1.**
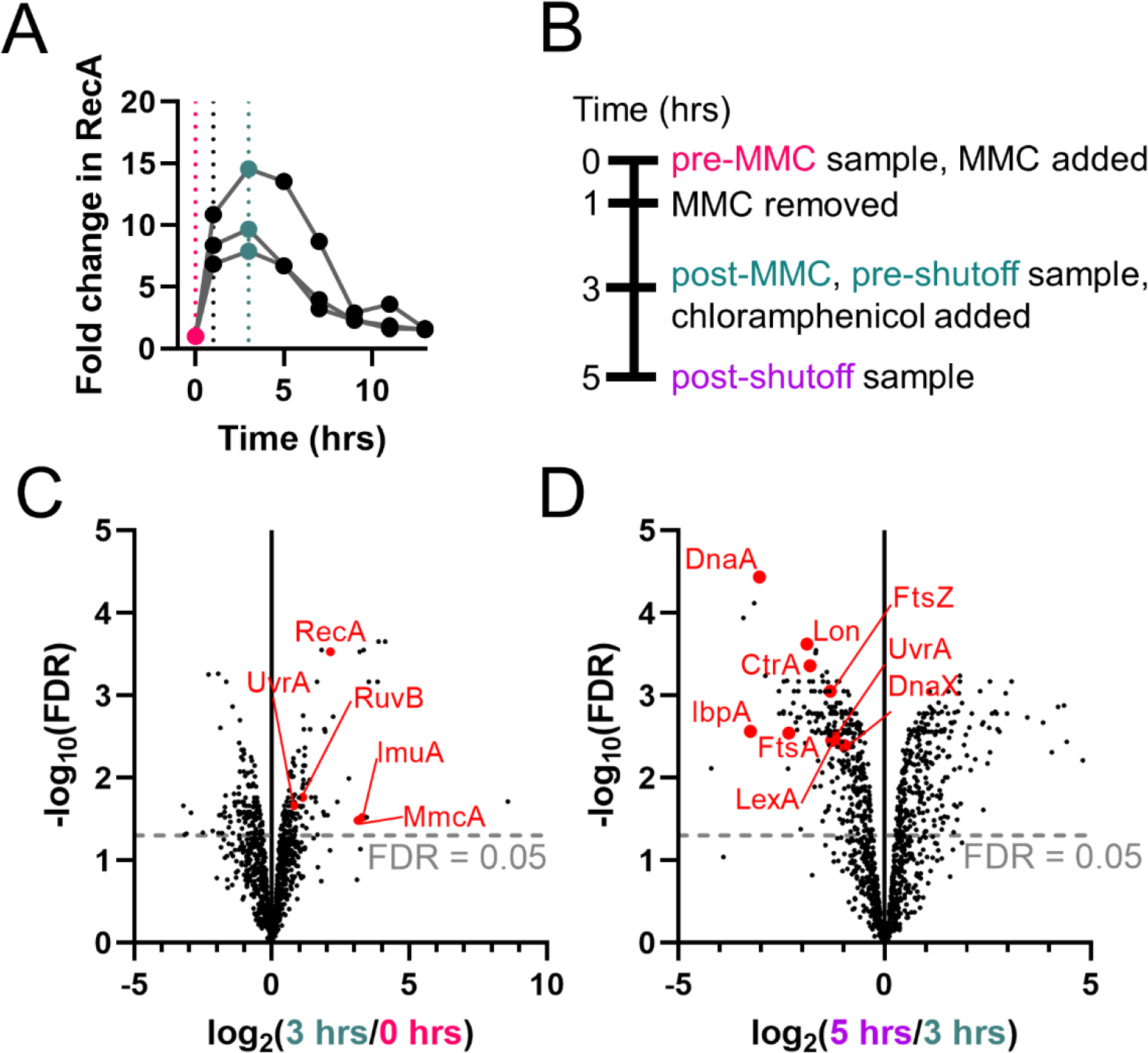
Survey of *Caulobacter* DNA damage response distinguishes known DNA damage response factors and protease substrates. (A) A western blot for RecA protein levels was used to determine peak protein induction (3 hours) to use for post-MMC timepoint for proteomics survey. Quantification of western blot for three replicates is shown, representative blot shown in Figure S1. RecA quantification was normalized to ClpP band and then to time 0 (Figure S1). (B) Samples for proteomics were taken before treatment (0 hours, pink line). Cells were treated with MMC for 1 hour and recovered in fresh medium for 2 additional hours before the post-treatment proteomics sample was taken (3 hours, teal line). At 3 hours chloramphenicol was added to shutoff translation and a post-shutoff sample was taken at 5 hours. (C) A volcano plot of proteomics data compares the statistical confidence (-log10(false discovery rate)) versus the change in protein abundance after MMC treatment log_2_(3 hrs/0 hrs). Several of the proteins that are significantly upregulated after DNA damage treatment are known DNA damage response proteins (highlighted in red). (D) A volcano plot of proteomics data compares the statistical confidence (-log10(false discovery rate)) versus the change in protein abundance after translation shutoff log_2_(5 hrs/3 hrs). Several of the proteins that are significantly downregulated after chloramphenicol treatment are known protease substrates (highlighted in red). FDR, false discovery rate.

This same timeline was used to obtain samples before and after MMC treatment for a proteomics survey (timescale in Figure 1B, results in Table S1, S2, and Figure 1C). In order to determine where post-translational regulation (e.g., proteolysis) may be affecting protein levels during DNA damage recovery, we treated cells with chloramphenicol at 3 hours to shut off translation and sampled cells for proteomics at 5 hours (timeline in Figure 1B, results in Figure 1C).

To validate our proteomics survey, we looked at the protein abundance changes of known DNA damage proteins after MMC treatment. As expected, several known DNA damage response proteins were more abundant after MMC treatment (e.g., RecA, UvrA, RuvB, ImuA, and MmcA, which are highlighted in red in Figure 1C). In the translation shutoff portion of the experiment, we observed a decrease in protein abundance of several known proteolysis substrates, including ClpXP substrates DnaX (25), CtrA (25), FtsZ (26), UvrA (13), and LexA(11), ClpAP substrates FtsA (26) and Lon (27), and Lon substrates DnaA (28) and IbpA (29) (all highlighted in red in Figure 1D). These data validated that our proteomics analysis could be used to identify known DNA damage response factors and proteolysis substrates.

Previous genome-wide surveys of the *Caulobacter* DNA damage response have focused on transcriptional regulation (18, 19, 22). We sought to determine how well the changes in protein abundance correlate to changes in RNA levels. To do this, we compared our proteomics data to previously published microarray data (19). In our western blot, we found that that RecA levels peak approximately 3 hours after MMC treatment and this data was consistent with a previous study (24). The same study indicated that RNA levels for *recA* and several other DNA damage response genes (e.g., *lexA, uvrA, ruvB, bapE*) peak at 60-80 minutes after MMC treatment. Based on this timeline and the consistency with our experimental design we used the study performed by Modell, et al. to compareour measurement of protein abundance changes to their published changes in RNA levels upon MMC treatment in PYE (19). In this study, transcript levels were monitored using microarrays over several time points (20-80 min) after MMC treatment (or hydroxyurea or ultra-violet light) in rich medium (or minimal medium) (19).

To discern between genes that are transcriptionally upregulated upon MMC treatment and those that are not, we used the maximum log_2_(Fold Change of RNA) from these timepoints for each gene and treated genes that were induced more than one standard deviation above the mean of these as “upregulated”. Genes that fell below this cutoff were considered “not transcriptionally upregulated.” This strategy highlighted 615 upregulated genes of which 146 could be identified as peptides in our proteomics survey. Of these, 43 show significant increase in protein abundance upon DNA damage (Table S4, log_2_(3 hrs/0 hrs) > 0, FDR < 0.05). As expected, the known DNA damage proteins mentioned above that showed increased protein abundances are included in this set (e.g., RecA, UvrA, RuvB, ImuA, and MmcA, which are highlighted in red in Figure 1C, all data in Table S4).

We identified 81 proteins that significantly increased in abundance upon DNA damage (log_2_(Fold Change in Protein) > 0, FDR < 0.05) that did not show a corresponding significant increase in RNA level by microarray as defined above ((19), Table S5). To further validate these hits, we compared them to a RNA-seq dataset that measured expression of genes following chronic MMC treatment (30). Of our list of 81 proteins, 76 showed no significant increase in expression after chronic MMC treatment (this study defined upregulated as a 3-fold increase in RNA levels after 4 hours (30)).

We followed up on three of the most induced proteins in this list to determine if they were important for survival to MMC, but deletion of these genes (CCNA_02118, CCNA_00591, and CCNA_00276; highlighted in purple in Figure 2A) had no effect on survival to MMC (Figure S2). These data indicate that while there are examples of proteins that are upregulated upon DNA damage that are not induced at the RNA level, not all of these are physiologically relevant to MMC survival.

**Figure 2.**
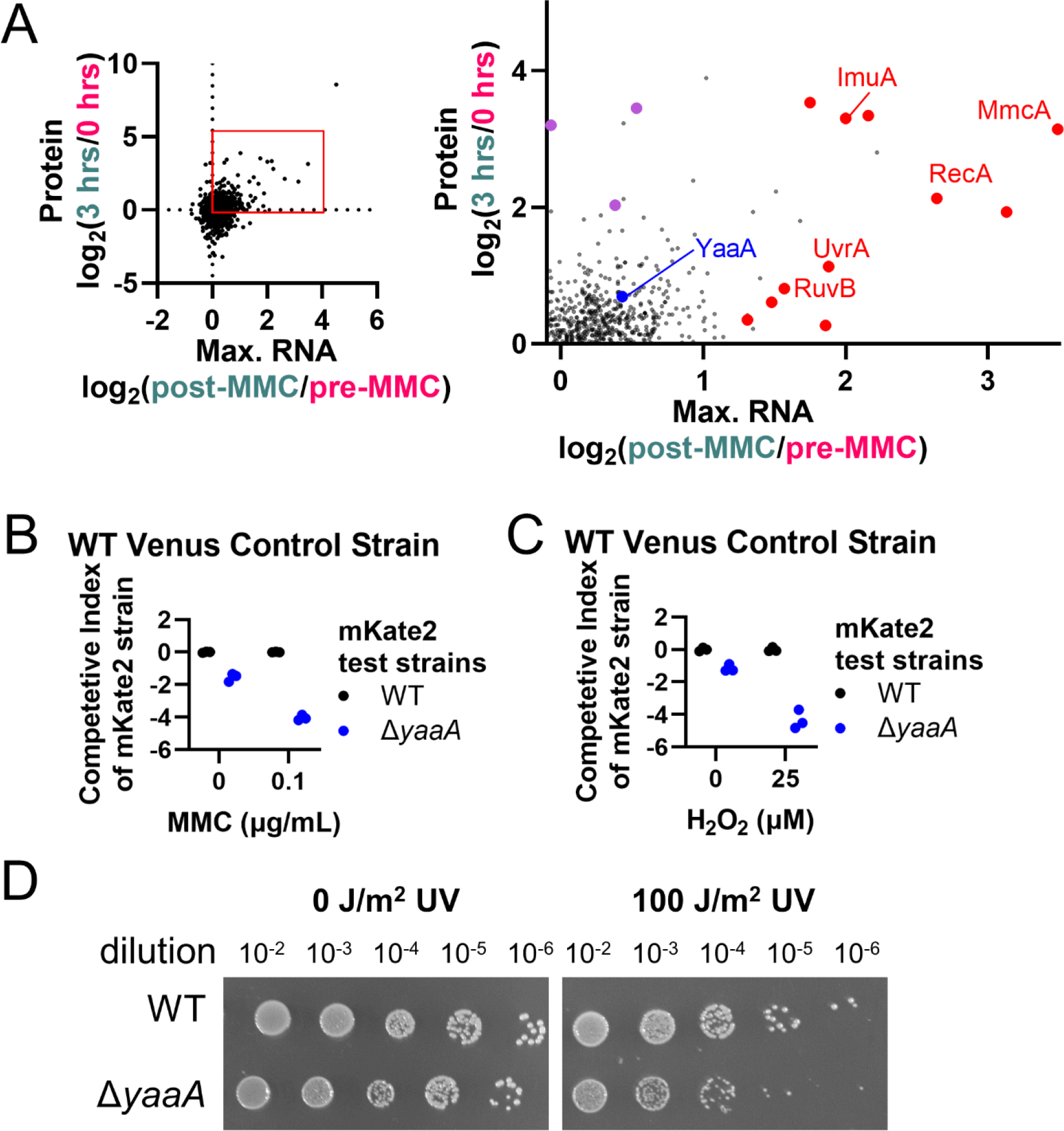
YaaA is upregulated upon MMC treatment and is important for survival to MMC, H_2_O_2_, and UV light. (A) A comparison between our proteomics and previously published microarray experiment that uses similar MMC treatment conditions (19). Graph on the right is an enlargement of the red box in the graph on the left. Proteins highlighted in red are known DNA damage response genes that are upregulated at the RNA and protein levels. YaaA is highlighted in blue. CCNA_02118, CCNA_00591, and CCNA_00276 are highlighted in purple. Graph cropped to upper right quadrant to highlight proteins that are upregulated. One gene is excluded (CCNA_03825) due to its extremely high induction to make other points more visible. (B and C) When each mKate2- expressing test strain is competed in co-culture against a wild type Venus-expressing fluorescent control strain, only the *ΔyaaA* mutant shows a competitive disadvantage that is exacerbated by the presence of (B) 0.1 μg/mL MMC or (C) 25 μM hydrogen peroxide. A reciprocal experiment in which test strains are marked with Venus and wild type control is marked with mKate2 is shown in Figure S3B and S3C. (D) The wild type and Δ*yaaA* strains were exposed to 0 or 100 J/m^2^ of ultra-violet light and 10-fold dilutions were plated. The Δ*yaaA* strain is more sensitive to ultra-violet light than the wild type strain.

Of this list of 81 proteins (Table S4) CCNA_03496 was also of particular interest. This gene is not upregulated upon MMC treatment in microarray (19) but does show a 17% increase in expression during chronic MMC treatment by RNA-seq (30). Due to its minor transcriptional induction upon DNA damage, YaaA had previously been overlooked as a potential DNA damage response factor. It has not previously been shown to be important for DNA damage survival and does not appear to be regulated by the LexA/RecA system (18, 22). CCNA_03496 is a homolog of *E. coli*’s *yaaA* ((31), we will refer to this gene as *yaaA* henceforth), which is important for regulating free iron levels during oxidative stress conditions (32). Our Tn-seq experiments identified it as potentially important for survival of MMC (Table S3) and published RB-Tn-seq results suggest that it is important for survival of the DNA damaging agent cisplatin (33). For these reasons, we decided to further characterize the connection between *yaaA* and MMC survival.

Deletion of *yaaA* had no effect on growth of isolated cultures treated with MMC (Figure S3A). We turned to more sensitive competition assays to quantify fitness defects. In these assays, we grew mKate2-expressing test strains with a Venus-expressing control strain and measured ratios of mKate2 to Venus fluorescence before and after several generations of growth. A Δ*yaaA* test strain expressing the mKate2 is less able than the wild type mKate2-expressing strain to compete in co-culture with the wild type Venus- expressing control (Figure 2C). This competitive disadvantage worsens in the presence of MMC (Figure 2B). To confirm that these results are not an artifact of the fluorescent strains, we reversed the markers to observe Venus-expressing test strains competed against a mKate2-expressing control and saw similar results (Figure S3B).

Because MMC activation can generate hydrogen peroxide (H_2_O_2_) in the cell (34), we sought to determine whether the sensitivity of the Δ*yaaA* strain to MMC was due to a sensitivity to DNA damage or to a secondary effect of possible H_2_O_2_ generation. We find that the Δ*yaaA* strain is indeed sensitive to H_2_O_2_ (Figure 2C and reciprocal experiment in Figure S3C). Importantly, the Δ*yaaA* strain is also sensitive to ultra-violet (UV) light, a non-chemical DNA damaging agent (Figure 2D). Taking these results together, the simplest explanation is that *yaaA* is important for survival to DNA damage resulting from photochemistry (UV), crosslinking/intercalation (MMC), or oxidative modification (H_2_O_2_).

We next looked at those genes downregulated following MMC treatment. As before, we considered a change in RNA levels of more than one standard deviation below the mean of all values as “downregulated.” Based on these criteria, we identified 87 proteins that became significantly less abundant upon DNA damage without a significant decrease in RNA abundance (Table S7). We performed further experiments on two genes that our Tn-seq results identified as more likely to contain transposon insertions upon MMC damage: *fzlC* and *purD*. However, deletion of these genes did not result in any change to MMC sensitivity (Supplemental Figure S4).

In the second stage of the proteomics survey, we compared protein level changes before and after translation shutoff (timeline in Figure 1B). We identified 320 proteins that significantly decreased in protein abundance (Table S8). Among these are the DNA damage response factors LexA and UvrA, which are known to be proteolyzed by ClpXP in *E. coli* (11, 13). These data suggest that this proteolysis may be conserved in *Caulobacter*. Others in this list may be substrates for proteolysis by ClpXP or other *Caulobacter* proteases.

These data suggest that while most protein abundance increases during the DNA damage response are explained by changes in gene expression, there are some exceptions. These exceptions may be instances where protein levels are controlled by post-transcriptional or post-translational regulation, but it is unclear how important these are to MMC survival. In our survey, we were able to identify and characterize a novel DNA damage response factor, YaaA, which is important to the survival of oxidative damage as well as the DNA damaging agents MMC and UV light. Finally, we found evidence to suggest that the proteolytic regulation of LexA and UvrA that was discovered in *E. coli* is conserved in *Caulobacter*.

### Investigation of the effect of *lon* deletion on the DNA damage response

The Lon protease is important for DNA damage survival in several bacterial species (16, 17, 35, 36), including *Caulobacter* (21). To investigate *lon’s* role in *Caulobacter’s* DNA damage response we performed the same proteomics experiment as above in a *Δlon* strain (full dataset in Table S1 and S2). Most of the known DNA damage-related proteins that were found to increase in abundance upon MMC treatment in the wild type strain also increase in the Δ*lon* strain (See RecA, RuvB, and MmcA in Figure 3A). The exceptions to this are UvrA, which did not increase in abundance upon MMC treatment in the *Δlon* strain (Figure 3A), and ImuA, which was not identified in the *Δlon* strain proteomics.

**Figure 3.**
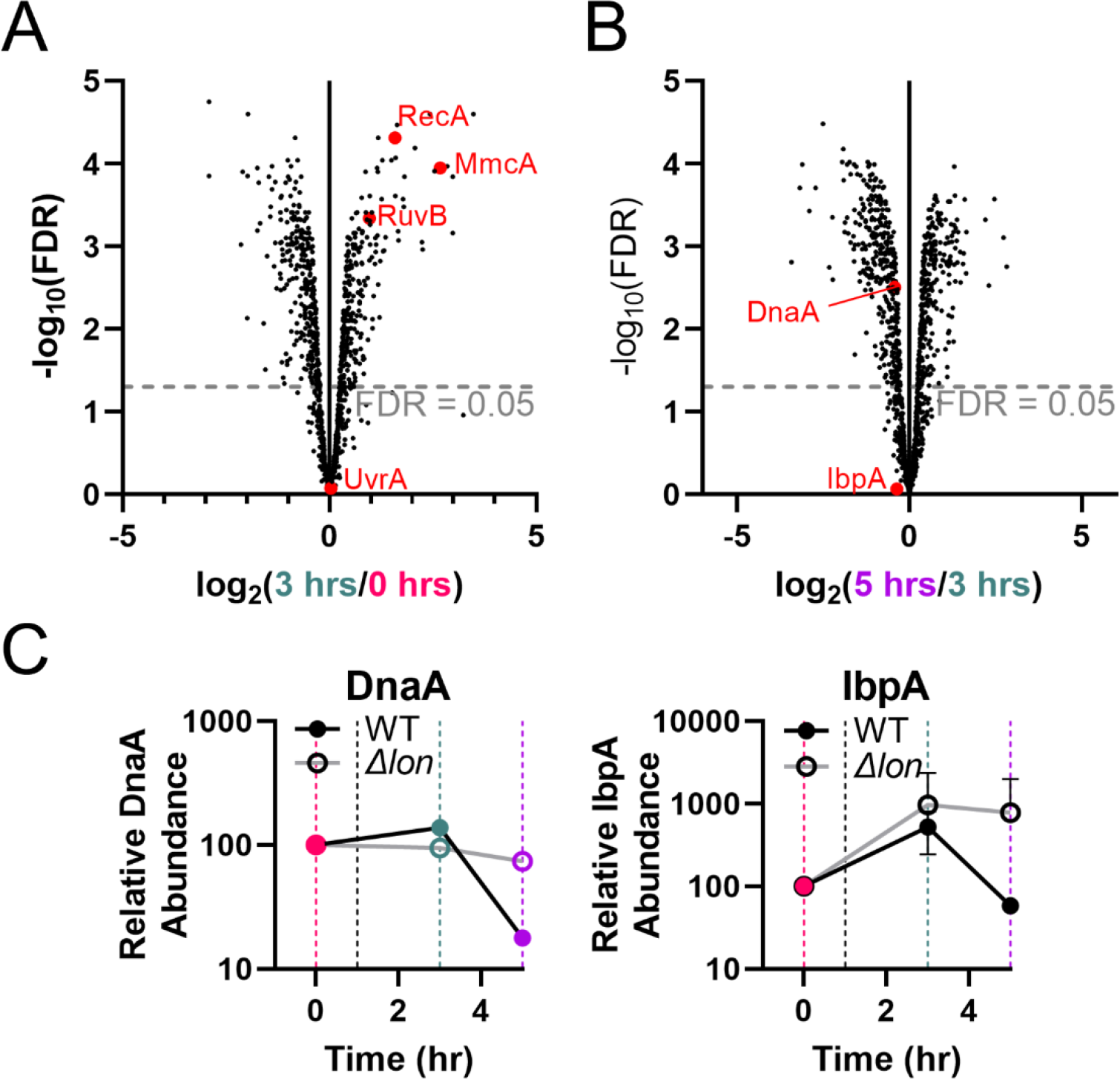
Proteomics Survey in *Δlon* strain distinguishes known Lon substrates. (A) A volcano plot of proteomics data compares the statistical confidence (-log10(false discovery rate)) versus the log_2_(3 hrs/0 hrs). Known DNA damage proteins shown in Figure 1B are highlighted here in red. ImuA was not identified in the proteomics of the Δ*lon* strain and so is absent. (B) A volcano plot compares the statistical confidence (- log10(false discovery rate)) versus the log_2_(5 hrs/3 hrs) for the Δ*lon* strain. Known Lon substrates DnaA and IbpA show a smaller decrease in protein abundance after translation shutoff than in the wild type strain (compare to Figure 1D). (C) Protein abundance of known Lon substrates DnaA and IpbA significantly increases upon MMC treatment (0-3 hours, pink and teal lines) and then decreases upon translation shutoff in a *lon*-dependent manner (3-5 hours, teal and purple lines). Note that protein abundances are not directly comparable between the two strains. FDR, false discovery rate.

In the translation shutoff portion of the proteomics experiment known Lon substrates DnaA and IbpA, which were shown to decrease in abundance in the wild type strain upon shutoff, showed a far smaller decrease in abundance in the *Δlon* strain (Figure 3B). As expected, decreases in protein abundance upon shutoff in the Δ*lon* strain are largely unchanged for ClpXP and ClpAP substrates (see examples for FtsZ, FtsA, and DnaX in Figure S5).

To identify potential Lon substrates from our list of potential protease substrates (Table S8), we compared the change in protein abundance after translational shutoff for each protein in the wild type and Δ*lon* strains. The 10% of proteins with the greatest difference in log_2_(5 hrs/3 hrs) between the two strains were considered potential Lon substrates (29 candidates; Table S8). The only known DNA damage factor identified in this list was UvrD; however, loss of *uvrD* is synthetic lethal with the deletion of *lon* (21, 37), suggesting that accumulation of UvrD is unlikely to underlie the Δ*lon* sensitivity to DNA damage. Overall, we consider it unlikely that stabilization of a single known DNA damage related protein is the primary driver for the increased sensitivity to DNA damage for the Δ*lon Caulobacter* strain. However, we did observe global changes to the DNA damage response in the Δ*lon* strain, which may be responsible for the MMC sensitivity.

Similar to the wild type strain, we monitored RecA levels through induction and recovery after MMC treatment in the Δ*lon* strain by western blot (Figure 4A and S1). RecA levels are about 3-fold higher in the Δ*lon* strain before MMC treatment (average RecA quantification normalized to ClpP quantification in wild type is 0.09±0.02 and in the Δ*lon* strain is 0.30±0.02, Figure S1). This is consistent with a previous observation that DNA damage response genes are upregulated in the Δ*lon* strain even in the absence of exogenous DNA damaging agents (23). In addition, peak RecA induction in the Δ*lon* strain occurs at a similar time point as the wild type (Figure 4A and S1) and both strains reach similar maximum RecA concentration at 3 hours (average RecA quantification normalized to ClpP quantification in wild type is 0.9±0.2 and in the Δ*lon* strain is 0.8±0.3, Figure S1). Together, this means that the fold-increase of RecA in the Δ*lon* strain is about 3-fold smaller of that observed in the wild type strain (Figure 4A and S1).

**Figure 4.**
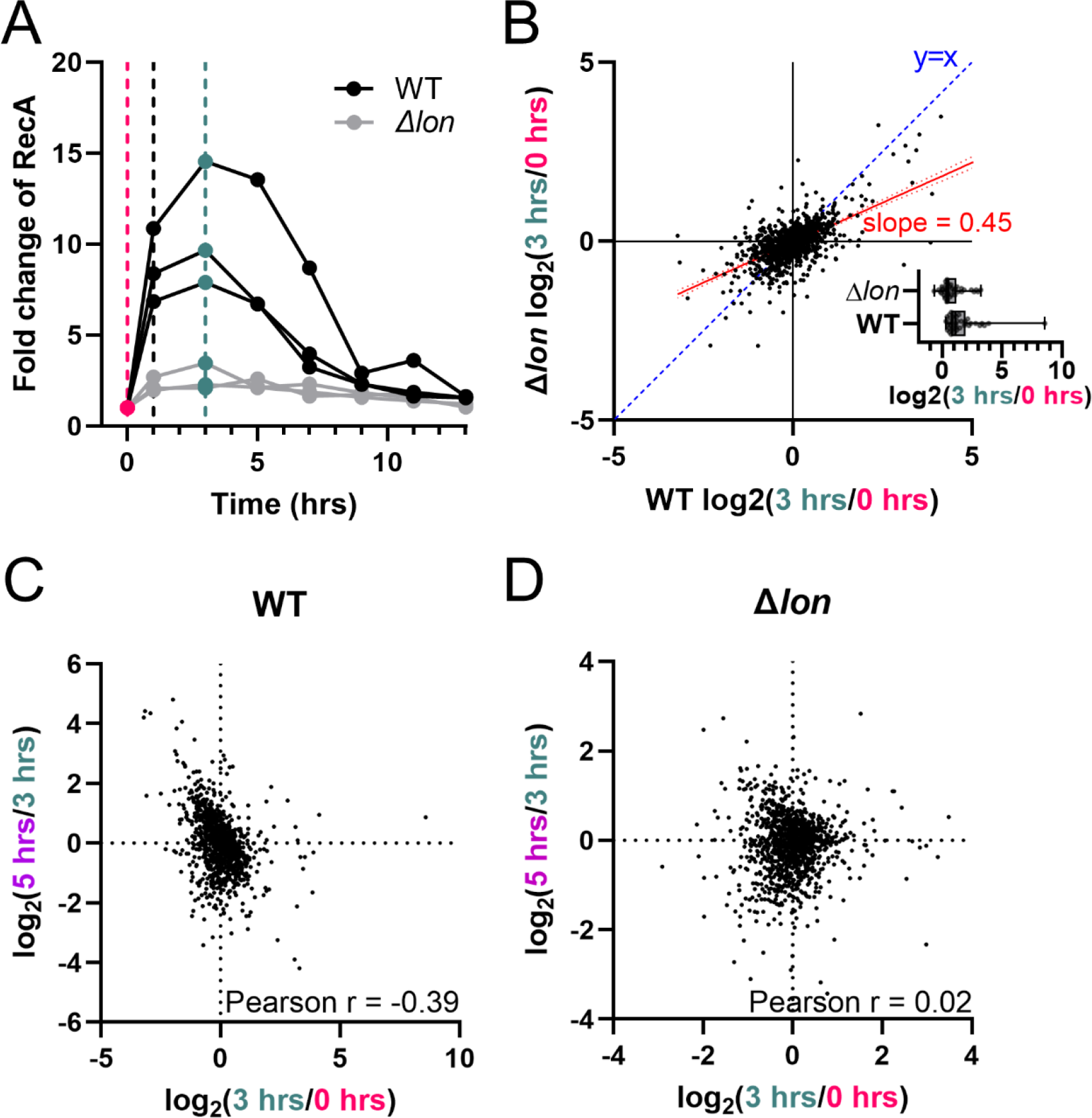
Protein abundance changes upon DNA damage are generally smaller in the Δ*lon* strain compared to in the wild type strain. (A) A western blot to determine RecA protein levels was used to determine when to sample for protein abundance changes after DNA damage treatment in the Δ*lon* strain. Note that the wild type data here is repeated from Figure 1A for easy comparison. Quantification of western blot for three replicates is shown, representative blot shown in Figure S1. Sampling for proteomics was performed as described in Figure 1B. RecA quantification was normalized to ClpP band and then to time 0 (see Figure S1). (B) Comparison of the log_2_(3 hrs/0 hrs) for the Δ*lon* versus wild type strain shows that fold changes of protein abundance are generally lower in the Δ*lon* strain. Blue dashed line represents an y=x line. The red line represents the trendline of the data with 95% CI drawn (slope = 0.45). The inset plots log_2_(3 hrs/0 hrs) for all proteins that are significantly upregulated (FDR < 0.05). Comparison of protein changes upon MMC treatment and translational shutoff indicates a negative correlation in the wild type strain (C) that is absent in the (D) Δ*lon* strain.

We found it to be a general trend that protein changes upon MMC treatment in Δ*lon* strain were less extreme than in the wild type strain (Figure 4B). Similarly, changes in protein abundance were dampened in general upon translation shutoff in Δ*lon* strain (Figure S6). Furthermore, previous studies have found evidence that highly expressed DNA damage response proteins are most rapidly degraded (38). We see a similar negative correlation between protein abundance changes upon MMC treatment (0-3 hours) and upon translational shutoff (3-5 hours) in the wild type strain (Figure 4C) that is absent in the Δ*lon* strain (Figure 4D). These proteome-level changes that we observe in the Δ*lon* strain may be related to *lon’s* importance to DNA damage survival in *Caulobacter*.

These data indicate that protein changes during the DNA damage response are generally explained by changes in RNA levels, but that there are some proteins that do not follow this trend. In our survey, we identified a novel DNA damage response factor, YaaA, which had been overlooked in previous transcription-focused surveys. The proteomics survey of the Δ*lon* strain was not able to identify a single critical Lon substrate that sufficiently explained the protease’s importance to MMC survival. However, we did observe global changes in protein dynamics that may be responsible for the MMC sensitivity of the strain.

## DISCUSSION

### Novel insights into the *Caulobacter* DNA damage response

By comparing our proteomics survey to previously published microarray data (19), we found that most protein abundance changes upon DNA damage are explained by changes at the RNA level, though there are exceptions (Figure 2A). Not all genes upregulated at the transcription or protein level were important for survival, which is consistent with the observations that genes upregulated during a stress response do not necessarily overlap with those genes important for tolerating that stress (39–41).

In addition, proteins encoded by three essential genes related DNA replication (*dnaQ, gyrA,* and *gyrB*) showed changes in protein abundance that are not explained by changes in RNA levels according to microarray (19) or RNA-seq (30). The *dnaQ* gene encodes the proofreading ε subunit of DNA polymerase III and its expression is upregulated upon MMC treatment in *E. coli* in a *recA*-dependent manner (42). Our proteomics survey suggests that its regulation may be post-transcriptional or post-translational in *Caulobacter*. Since its protein abundance decreases after translational shutoff (Table S2), proteolysis may play a role. Levels of another replisome component, DnaX, is also regulated differently in *E. coli* and *Caulobacter.* In *E. coli* a truncated form of DnaX (γ) that is important during DNA damage (43) is created through a ribosomal frameshift (44), while in *Caulobacter* similar DNA damage-related γ forms (45) are created through partial proteolysis by ClpXP (46). It is possible that proteolysis regulates the concentrations of both of these replisome components in *Caulobacter*.

We find that DNA gyrase levels (subunits A and B) decrease upon MMC treatment in *Caulobacter* (Table S2) but RNA levels do not appear to decrease when measured by microarray (19, 22) or RNA-seq (30). DNA gyrase levels continue to decrease after translational shutoff, indicating that the abundance of this protein may be regulated by proteolysis (Table S2). In *E. coli* overexpression of a chromosomally encoded DNA gyrase inhibitor *gyrI* increases survival to MMC (47, 48). It’s possible that *Caulobacter*’s downregulation of DNA gyrase is filling a similar role.

We also identified a novel DNA damage response factor, YaaA, which has previously been shown to be important for survival to oxidative damage in *E. coli* (32). In *Caulobacter*, we show that *yaaA* is important to the survival of the DNA damaging agents MMC and UV light (Figure 2B-2D), indicating that its physiological relevance is not limited to oxidative damaging conditions. It’s unclear if *yaaA’s* phenotypes in *Caulobacter* are related to its role in regulating iron levels as seen in *E. coli* (32).

Previous work has suggested a correlation between level of expression during the DNA damage response and rate of proteolysis (38). This proteomics survey supports this model, as we see a negative correlation between protein abundance changes upon MMC treatment and those upon translation shutoff (Figure 4C). Several DNA damage repair proteins have been identified as proteolysis substrates in *E. coli,* including the LexA N-terminal domain, UvrA, RecA, RuvB, and RecN (38). Of these, our proteomics data provides evidence to support this categorization for the LexA N-terminal domain, UvrA, and RecA but RuvB and RecN do not decrease in protein abundance upon translation shutoff in our survey (Table S2).

This proteomics survey has identified examples of proteins that are regulated independently of transcription during the DNA damage response, though it is unclear how many of these are physiologically relevant to the response. We have also identified instances where regulation of certain proteins during DNA damage in *E. coli* appears to be conserved in *Caulobacter*, however the mechanisms of regulation appear to differ in some of these cases. Finally, we have identified a novel DNA damage response factor that has been overlooked in previous surveys that have focused on transcription alone. This highlights the utility of combining transcriptomic and proteomic approaches to screen for novel factors.

### The Lon protease and the DNA damage response

Since DNA damage related Lon substrates UmuD (14) and SulA (15) discovered in *E. coli* are not conserved in *Caulobacter*, it is unclear why *lon* is important to MMC survival (21). We attempted to identify novel Lon substrates but did not find any potential substrates that sufficiently explain this phenotype (Table S8).

The Δ*lon* strain exhibits global differences from the wild type strain in protein dynamics that may account for the strain’s sensitivity to MMC (Figure 4 and S6). We find that protein abundance changes during the DNA damage response are dampened in the Δ*lon* strain (Figure 4B and S6), which is at least partially due to the higher basal level expression of DNA damage response genes in the strain (23). We also find that the correlation between changes in protein abundance upon DNA damage treatment and those upon translation shutoff is absent in the Δ*lon* strain (Figure 4C and 4D). These proteome-wide changes to the DNA damage response may be at least partially responsible for the strain’s sensitivity to MMC, possibly due to a diminished range of response dynamics.

### Concluding thoughts

In our proteomics analysis, we identified and quantified protein abundances for about 30% of the *Caulobacter* proteome. While some of the missing proteins are likely not expressed during the growth conditions of our assays, some key DNA damage response proteins are missing from our dataset. Some of these are known to be membrane-associated (e.g. SidA (19) and DidA (22)), which may reduce our detection, and others likely have relatively low abundance, even during DNA damage induction. Future work using protocols designed specifically to observe these proteins could reveal additional novel factors involved in DNA damage survival or reveal novel mechanisms of regulation for known factors.

Our survey of protein dynamics during the DNA damage response allowed us to confirm that, in general, protein abundance changes during the response are explained by changes in RNA level. We also cataloged exceptions to this general rule and characterize the novel DNA damage response factor, YaaA. Finally, though we did not identify novel Lon substrates that explain this protease’s importance to DNA damage survival, we did identify systems-level differences in the size of protein abundance changes during the response in the absence of *lon* which may play a role in this phenotype.

## METHODS

### Bacterial strains and growth conditions

Bacterial strains and plasmids used in this work are listed in Tables S9 and S10, respectively. All *Caulobacter* strains used in this work are NA1000 derivatives. *Caulobacter* strains were grown in PYE broth (2 g/L peptone, 1 g/L yeast extract, 1 mM MgSO_4_, and 0.5 mM CaCl_2_). Solid media was made with 1.5% agar. Knockout strains were created by replacing the native gene with a gentamycin resistance cassette using the pNPTS138 non-replicating vector (45). Venus- and mKate2-expressing strains were created by integrating pXGFPC-2 (49) based plasmids encoding constitutively expressed fluorescent proteins driven by a *lac* promoter (50) at the xylose locus.

### Bacterial growth curves

An overnight culture was diluted into PYE at an OD600 = 0.05. A volume of 200 μL of cells was added to wells of a 96-well plate. Plate was shaken continuously at 30°C and OD600 was read every 20 minutes in a BioTek Epoch 2 Microplate Spectrophotometer (Agilent, Santa Clara, CA). MMC (0.5 μg/ml; Millipore Sigma, St. Louis, MO) was used to treat cells with DNA damage.

### Western blot

Western blots were performed using a commercially available rabbit anti-*E. coli* RecA primary antibody (1:10,000 dilution; Abcam, Cambridge, MA, 16 hours). Blots were visualized using the goat anti-rabbit Alexa Fluor™ Plus 800 secondary antibody (1:10,000 dilution; Licor, Lincoln, NE, 1 hour). Bands were quantified using ImageJ (51).

### Competition assays

Overnight cultures of each test strain and the control strain were diluted to an OD600 = 1.0, mixed 1:1, and serially diluted 30,000-fold. Undiluted cells were analyzed with a SpectraMax M2 microplate reader (Molecular Devices, San Jose, CA). Measurements were taken for cell density (OD600), Venus fluorescence (Excitation 490 nm, Cut off 515 nm, Emission 530 nm), and mKate2 fluorescence (Excitation 575 nm, Cut off 590 nm, Emission 630 nm). Diluted samples were grown to stationary phase (∼48 hours, shaking at 30ᵒC) and optical density and fluorescence measurements were taken again. The competitive index was calculated as the log_2_[((ratio test/control)END / (ratio test/control))START / ((average ratio wild type test/control)END/[(ratio wild type test/control))START].

### Sensitivity testing to ultra-violet light

Cells were grown to exponential phase and treated with 0 or 100 J/m^2^ with a UV Stratalinker (Strategene, San Diego, CA, USA). Cells were diluted 10^0^-10^-7^ and dilutions were plated on PYE agar. Plates were incubated for 48 hours at 30ᵒC and then photographed.

### Transposon Sequencing

Transposon libraries were created as previously described (52, 53). A wild-type EZ-Tn5 library was treated in triplicate with mitomycin C (0.5 μg/mL) for 1 hour, outgrown for about 10 generations, and sequenced. Data was analyzed using edgeR (54). Differential analysis of the counts in these libraries is available in Table S3.

### Sample collection and preparation for proteomics

#### Pre-damage sample

Three biological replicates of overnight growth of each *Caulobacter* strain (wild type and *Δlon)* were diluted to an OD600 = 0.0005 using PYE to a volume of 50 mL and grown for approximately 10 generations to an OD600 of 0.5-0.6. All samples were normalized to an OD600 of 0.5 using PYE to a volume of 25mL. A 5 mL pre-damage sample was taken, and cells were collected by centrifugation (6,000 x g, 5min). Cell pellets were resuspended in fresh lysis buffer (8 M urea, 50 mM HEPES pH7.5, 75 mM NaCl), flash frozen, and stored at -80°C.

#### Post-damage/pre-shutoff sample

Cells were treated with 0.5 μg/mL MMC for 1 hour before cells were collected by centrifugation (6,000 x g, 10 min) and resuspended in fresh PYE broth to an OD600 = 0.5 to a volume of 25 mL. These treatment conditions are similar to those of the microarray from the Laub lab (19) for simple comparison. Previous experiments indicate that the wild type strain has about 55% survival to this treatment (45). After 2 hours, all samples were normalized to an OD600 = 0.5 using PYE to a volume of 25 mL. A 5 mL post-damage sample was taken, and cells were collected by centrifugation (6,000 x g, 5 min). Cell pellets were resuspended in fresh lysis buffer, flash frozen, and stored at -80ᵒC.

#### Post-shutoff sample

Cells were treated with chloramphenicol (30 μg**/**mL) for 2 hours. A 5 mL post-shutoff sample was taken, and cells were collected by centrifugation (6,000xg, 5 min). Cell pellets were resuspended in fresh lysis buffer, flash frozen, and stored at -80°C.

#### Sample preparation for proteomics

Samples were freeze-thawed three times in liquid nitrogen to aid cell lysis, then centrifuged at 16,000 x g for 10 min at 4ᵒC. Supernatant was carefully transferred to a fresh 1.5mL tube. A Bradford assay was performed to quantify protein concentration. Because the Bradford assay is sensitive to high urea concentrations, a small fraction of each sample was diluted 4-fold for the Bradford assay. 50 μg of each sample was transferred to a fresh 1.5mL tube. The final volume was adjusted to 50 μL using 100 mM Triethylammonium bicarbonate (TEAB) buffer.

To reduce disulfide bonds, 5 μL 200 mM tris(2-carboxyethyl)phosphine (TCEP) was added to each sample and incubated at 55°C for 1 hour. To modify cysteine residues, 5 μL of 375 mM iodoacetamide was added to each sample and incubated 30 min at room temperature in the dark. To precipitate protein, 6-volumes of cold acetone was added to each sample, and they were stored overnight at -20°C.

Samples were centrifuged at 8,000 x g for 10 min at 4°C to collect precipitate. Supernatant was carefully removed and discarded. Pellets were allowed to dry at room temperature for 1 hour. Protein pelleted were resuspended in 100 μL 50 mM TEAB. Samples were incubated overnight at 37°C with 1.25 μg of trypsin (for a 2.5:100 ratio of trypsin to protein).

### Tandem mass tagging for proteomics

Samples were labelled using the TMT10plex™ Isobaric Label Reagent Set (ThermoFisher Scientific, Waltham, MA, Catalog #90110) according to the product instructions. Since each sample only contains 50 μg of peptides instead of the suggested 100 μg, half of each label was used per sample.

### Proteomics sample fractionation

For each replicate experiment in each strain, equal volumes of each samples were combined into a single tube and then dried in a SpeedVac. Pooled samples were resuspended in 300 μL 0.1% TFA. Samples were dried in a SpeedVac a second time and resuspended in 300 μL 0.1% TFA. The pH of each sample was measured to confirm that it was between 2.5 - 3.5, which is critical for effective fractionation.

Pooled samples were fractionated using the Pierce™ High pH Reversed-Phase Peptide Fractionation Kit (ThermoFisher Scientific, Waltham, MA, Catalog #84868) according to the kit instructions. Fractions 1 and 5, 2 and 6, 3 and 7, as well as 4 and 8 were pooled together for a final total of 4 fractions per sample. Samples were dried and resuspended in 0.1% folic acid in water for mass spectrometry.

### Mass spectrometry for tandem mass tagging proteomics

LC-MS analysis was performed, modified from (55) using an Easy-nLC 1000 nanoLC chromatography system interfaced to an Orbitrap Fusion mass spectrometer (ThermoFisher Scientific, Waltham, MA). Samples were pre-concentrated and desalted on to a C18 trap column prior to separation over a 180 min gradient from 0% to 50% mobile phase B (A:0.1% formic acid in water, B:0.1% formic acid in acetonitrile) at 300 nL/min flow rate with a 75 µm x 15 cm PepMap RLSC column (ThermoFisher Scientific, Waltham, MA). Mass spectrometry parameters were as follows: ion spray voltage 2000V, survey scan MS1 120k resolution with a 2 s cycle time, interleaved with data-dependent ion trap MS/MS of highest intensity ions with charge state-dependent ion selection window (z=2:1.2 m/z, z=3:0.7 m/z, z=4-6:0.5 m/z) and CID at 35% normalized collision energy. Additionally, the top 5 (z=2) or 10 (z>2) product ions were synchronously selected for HCD MS^3 at NCE 65% with Orbitrap detection to generate TMT tag intensities.

### Proteomics Data Analysis

RAW data files were analyzed in Proteome Discoverer 2.4 (ThermoFisher Scientific, Waltham, MA) using the SEQUEST search algorithm against *Caulobacter crescentus* NA1000 (NCBI Reference Sequence: NC_011916.1) database downloaded from uniprot.org. The search parameters used are as follows: 10 ppm precursor ion tolerance and 0.4 Da fragment ion tolerance; up to two missed cleavages were allowed; dynamic modifications of methionine oxidation and N-terminal acetylation. Peptide matches were filtered to a protein false discovery rate of 5% using the Percolator algorithm. Peptides were assembled into proteins using maximum parsimony and only unique and razor peptides were retained for subsequent analysis. Protein quantitation based on TMT ion abundance was performed using a co-isolation threshold of 75% and SPS match threshold 65%. Each TMT channel was normalized to total peptide amount and then abundance scaled to average 100% for all proteins.

Protein abundances were then normalized to the averages of the 0 hour samples (Table S1). Protein ratios (Table S2) were calculated using edgeR (54).

## ACKNOWLEDGEMENTS

This work was supported by NIH/NIGMS R35GM130320 to P. Chien. The mass spectrometry in this paper was performed by Dr. Stephen Eyles at the University of Massachusetts Amherst Mass Spectrometry Core Facility in Amherst, MA (RRID:SCR_019063). The *ΔfzlC* strain was a generous gift from Erin Goley’s lab at Johns Hopkins University, Baltimore, MD.

