## Supplemental Figures 1-6 for "Proteomic survey of the DNA damage response in *Caulobacter crescentus*"

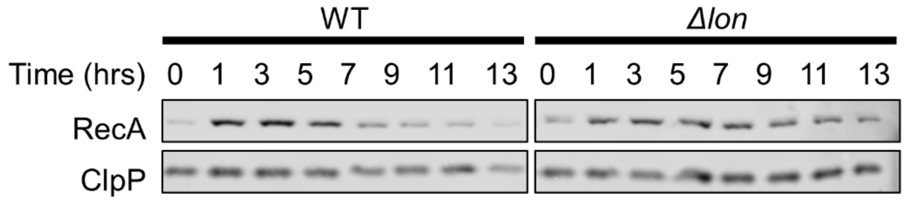


**Figure S1. Western blot for RecA reveals timeline of DNA damage induction and return to baseline.** Cells were grown to exponential phase and treated with 0.5 μg/mL MMC for 1 hours (0-1 hours) before MMC was removed. At each timepoint, cells were diluted to an OD600 of 0.5 to maintain exponential phase growth. Experiment performed in triplicate, one representative image shown. Quantification of wild type blots are shown in Figure 1A. Quantification of Δ*lon* blots are shown in Figure 4A.

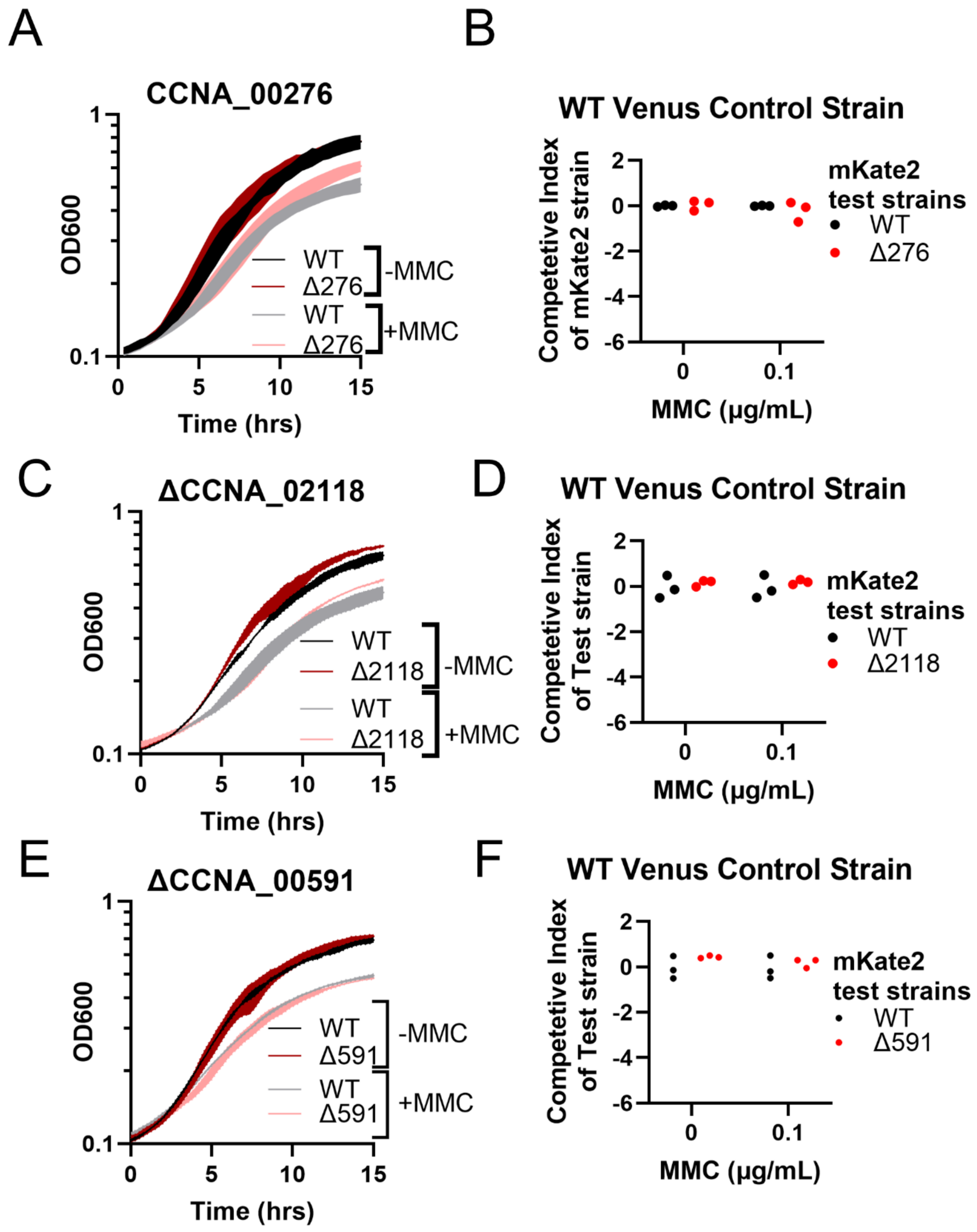


**Figure S2. Deletion of CCNA_00276, CCNA_02118, or CCNA_00591 had no effect on MMC sensitivity.** Growth curves (A, C, E) and competition assays (B, D, F) show that deletion of CCNA_00276 (Δ276, A and B), CCNA_02118 (Δ2118, C and D), or CCNA_00591 (Δ591, E and F) does not affect MMC sensitivity.

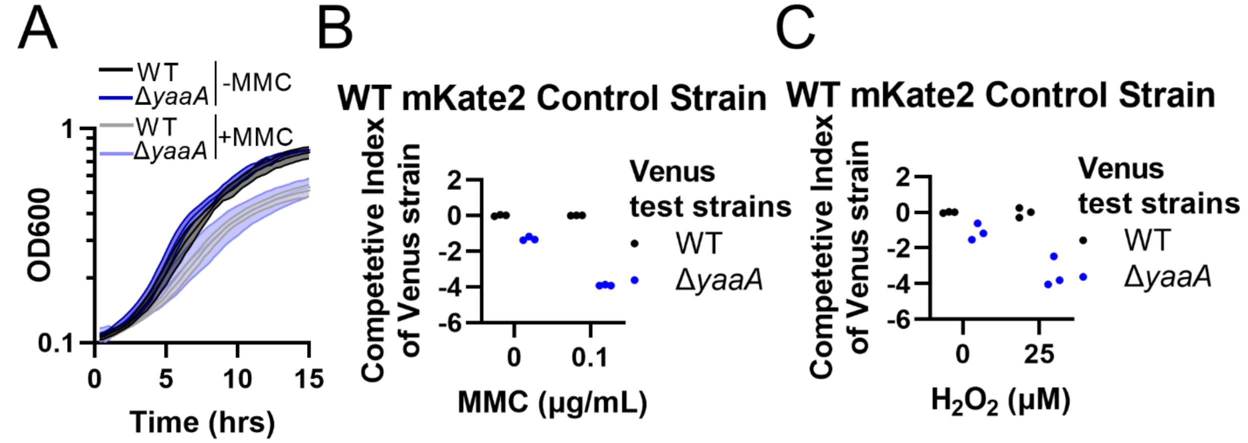


**Figure S3. Supplemental data corresponding to Figure 2 in the main text.** (A) Growth curves of Δ*yaaA* in 0 or 0.5 μg/mL MMC show no significant differences in growth when compared to the wild type strain. When each Venus-expressing test strain is competed against in co-culture a wild type mKate2-expressing fluorescent control strain, only the Δ*yaaA* mutant shows a competitive disadvantage that is exacerbated by the presence of (B) 0.1 μg/mL MMC or (C) 25 μM hydrogen peroxide. Reciprocal experiment where test strains are marked with mKate2 and wild type control is marked with Venus is shown in Figure 2C in main text.

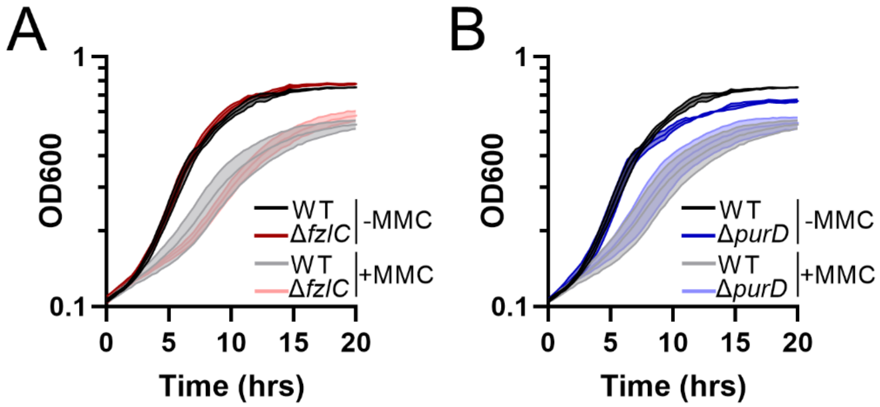


**Figure S4. Deletion of two genes encoding proteins that are downregulated upon DNA damage does not affect MMC survival.** (A) The Δ*fzlC* strain show no difference in MMC sensitivity compared to the wild type strain. (B) The Δ*purD* strain has a stationary phase defect that is not observed in the presence of MMC. All experiments were performed in triplicate. Mean and standard deviation are shown. Same data for wild type controls are shown in (A) and (B) for ease of comparison.

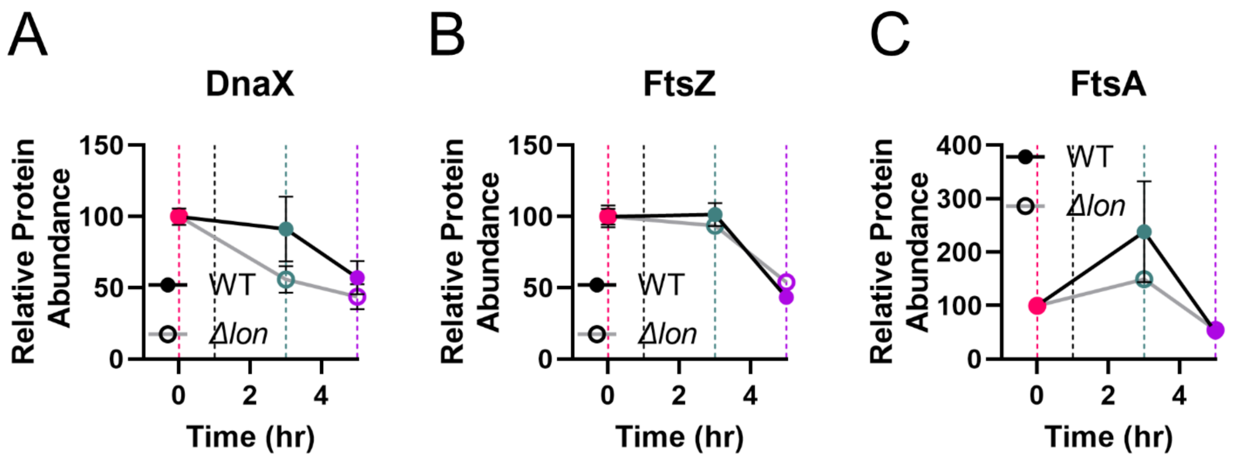


**Figure S5. Protein abundance changes in known ClpXP and ClpAP substrates.** Proteomics results for changes in DnaX, FtsZ, and FstA protein abundance upon MMC treatment and subsequent translational shutoff. Samples for proteomics were taken before treatment (0 hours, pink line). Cells were treated with MMC for 1 hour and recovered in fresh PYE for 2 additional hours before the post-treatment proteomics sample was taken (3 hours, teal line). At 3 hours, cells were treated with chloramphenicol and final proteomics samples were taken after 2 additional hours (5 hours, purple line). Note that protein abundances are not directly comparable between the two strains.

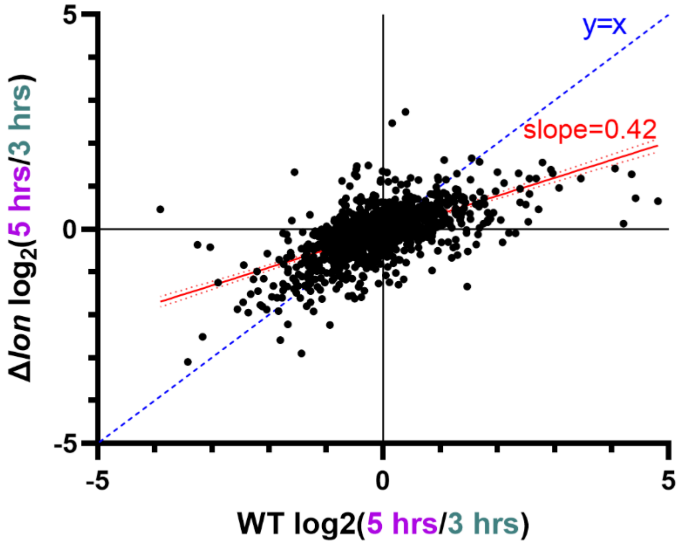


**Figure S6. Proteolysis upon translational shutoff is overall lower in the Δ*lon* strain.** Comparison of the $\log_2$(fold change in protein abundance upon translational shutoff) for the Δ*lon* versus wild type strain shows that fold changes of protein abundance are generally lower in the Δ*lon* strain. Blue dashed line represents an y=x line. The red line represents the actual trendline of the data with 95% CI drawn (slope = 0.42).
